# Mental Schema Reduces Cognitive Load and Facilitates Emergence of Novel Responses in Mice and Artificial Neural Networks

**DOI:** 10.1101/2024.10.23.619968

**Authors:** Vikram Pal Singh, Shruti Shridhar, Shankanava Kundu, Richa Bhatt, Balaji Jayaprakash

## Abstract

Mammalian brain has evolved to infer from past experiences and elicit context relevant novel behavioural responses hitherto unexpressed by the animal. However, little is known about how prior knowledge influences the emergence of such responses. Remarkably, the brain not only arrives at these responses through logical inferences based on previous leanings, but also acquire new related information, without causing catastrophic interference. Mental schemas have often been proposed as the framework for this phenomenon. In this study, using mice as a model animal, we show that schematic networks not only enhance the cognitive load handling capacity (CLHC) and prevent catastrophic interference, but also facilitate the generation of novel, contextually relevant responses. Interestingly, when the animals were trained in a paradigm that did not invoke the pre-formed mental schema, we observed neither an enhancement to CLHC nor a generation of novel context relevant responses. Based on the principles of mental schemas discovered in our animal experiments, we developed a biologically plausible artificial neural network (ANN) that avoids catastrophic interference and captures the learning properties observed in our experiments. The custom architecture of this ANN enables it to generate responses similar to those of animals in novel scenarios.

**Significance Statement:** Little is known about the role of mental schemas in preventing memory interference—a process in which overlapping or similar memories hinder the acquisition and retention of related information. In this study, we demonstrate that mental schemas enhance cognitive load handling capacity and improve the ability to solve novel but related problems. Using mice as a model, we show that the mere existence of a mental schema is not enough for improved cognitive load handling; instead, the relationship between existing and new information must be explicitly established during the learning process. Based on these findings, we developed a minimalistic artificial neural network (ANN) that effectively mimics this behaviour. These insights pave the way for developing more efficient learning and teaching strategies.

## Introduction

The organization of information as knowledge structures in the brain (*mental schemas*) has been widely postulated and studied in humans (Bartlett 1932; Carmichael, Hogan, and Walter 1932). Bartlett (Bartlett 1932) hypothesised that storing the past experiences in organised knowledge structures such as mental schema aids in forming new ideas through “*constructive and predictive processes”*. While a few studies have investigated the existence of mental schemas and their role in system consolidation across various animal species (Farovik et al. 2015; Mao et al. 2021; McKenzie et al. 2014; Tse et al. 2007),these have typically focused on either the role of mental schemas in rapid acquisition and consolidation or on the brain regions associated with them. The precise role of schemas in complex cognitive phenomena, such as problem-solving (Benedek et al. 2023; Lebuda and Benedek 2023) and managing higher cognitive loads during learning, remains largely unexplored, especially in animal models. In our previous work, we adapted a mental schema task from rats (Day, Langston, and Morris 2003) to study temporal order memory in mice (Shridhar et al. 2022). Having successfully translated this task to mice in our lab, we now use the same setup with a new experimental design to investigate the role of mental schemas in preventing catastrophic forgetting and facilitating problem-solving.

The conventional approach to studying learning and memory primarily focuses on understanding associations formed through stimuli-stimuli or stimulus-response presentations (Heath 1979; Pearce 1987; Pavlov 2010; R. A. Rescorla 1972). While these methods help uncover fundamental principles of learning, they are limited in explaining the abstraction of interstimulus relations that emerge when multiple equally contingent associative pairs are presented simultaneously (Asch 1969). We hypothesized that a framework like mental schema could address this limitation by expanding cognitive load handling capacity (CLHC) and providing a more efficient learning system that integrates pre-learned knowledge. Such a system could balance storing new information while relating it to past knowledge (van Kesteren et al. 2012). We propose that mental schemas, as organized structures, serve as templates against which new information can be compared and stored, thereby enhancing CLHC. Specifically, we hypothesize and test whether prior knowledge in the form of schemas facilitates the acquisition of complex information that would otherwise be difficult to learn. To test this, we designed an event arena-based flavour-place association task for mice.

## Results

### Event Arena Based Flavour Place Association Paradigm in Mice

Our experimental design involves training mice in a flavour-place association task within an event arena (*Fig. 1a*; see Methods for full details). The event arena consists of a cuboidal sandbox with 5 sand wells. In this task, the animals are required to find the correct location (sand well) containing the food pellet. They are given 120 seconds to explore the arena and retrieve the pellet from the correct well. The goal is for the animals to learn the association between each flavour and its corresponding location. Upon entering the arena, the animals explore and learn to associate the location of the reward sand well with the flavour presented at the start of the trial (see methods and Shruthi et al 2022). The locations of the sand wells are selected to minimise rotational symmetry. The arena has 2 distinct intra-maze cues that defines the context. Training consists of multiple sessions. Each session consists of 5 trials, one for each flavour-place pair. During training, the animals must learn 5 distinct flavours and their corresponding locations in the arena. The flavour-place pair is kept unique and consistent throughout the experiment. The sessions are spaced at least 2 days apart and trials are separated by 45-60 mins. Both the entry start box and the order of flavour presentation are randomized, while maintaining consistent flavour-to-location correspondence across all trials and sessions.

**Fig 1:**
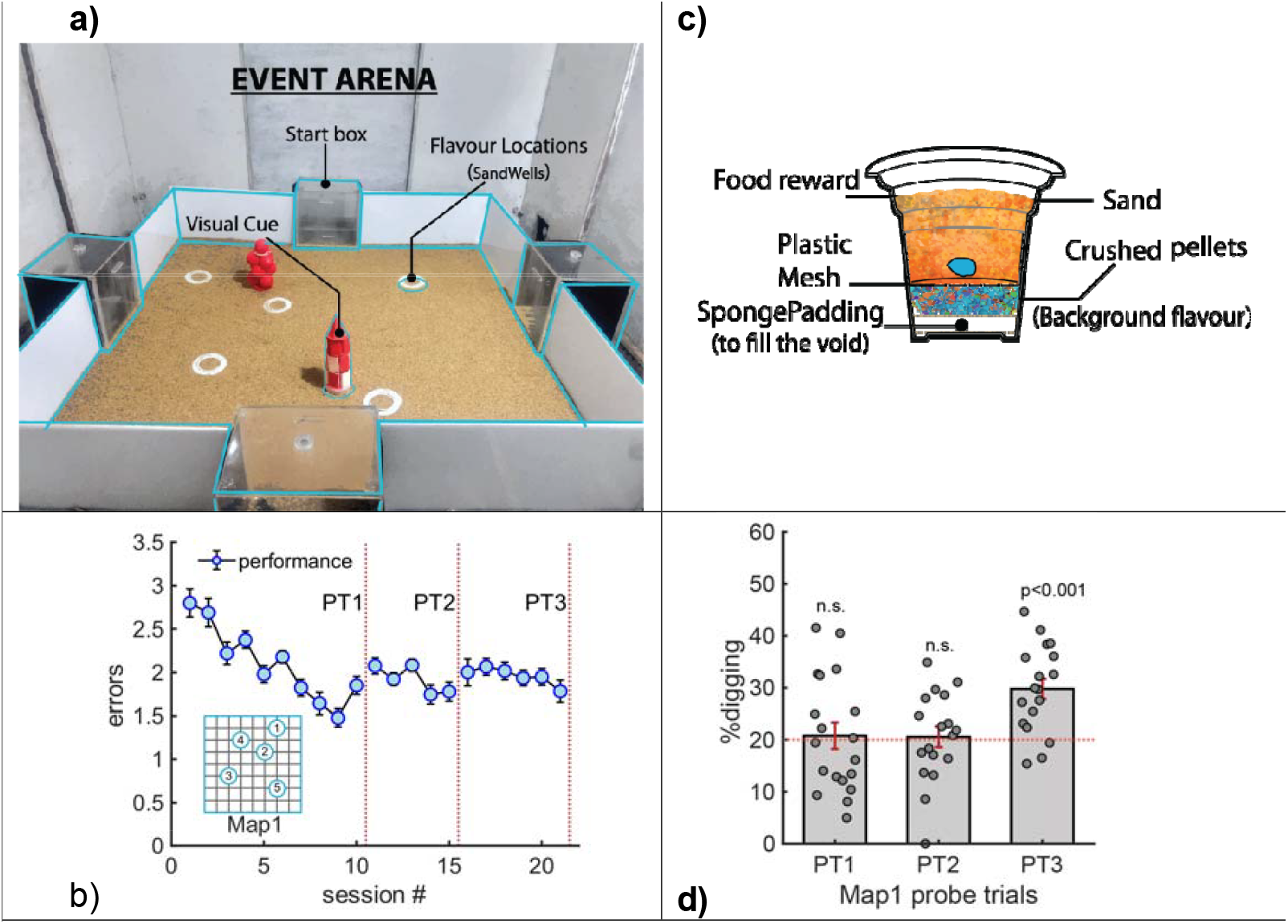
Multiple flavour-place association paradigm in mice. **(a)** Event Arena (1.04 m x 1.04 m) with 5 sand wells accessible at any given point from the start boxes decided by the experimenter. **(c)** The custom designed sand wells with the base filled with crushed flavour pellets excluding the flavour on the mesh so as to produce same background odour across all the sand wells. **(b)** Acquisition of paired associates over a period of 21 sessions. Red dashed lines indicate the sessions corresponding to probe trials. The inset illustrates the configuration of flavour locations for Map#1. **(d)** The performance of the animals improves over time in a cued recall probe trial with performance significantly above chance after session 21 in PT3 (n=18, One Sample t-test, t-Statistics 4.91525, p=1.308 x 10 ^-4^).

Animal performance is recorded in terms of errors, defined as the number of incorrect locations scratched or dug during a trial. We observed that the number of errors decreased progressively across sessions (*Fig1b*). A key measure of learning is the ability to recall, which is typically assessed by evaluating the persistence of correct responses during a probe trial (cued but unrewarded). On the day of the probe trial, animals are cued with a flavoured pellet outside the arena (as in training), but there is no accessible food reward at the correct location. To prevent the animals from being guided by odour cues, food pellets corresponding to all 5 flavours are placed beneath a mesh in each well (*Fig1c*; See methods for complete description of flavour pellet preparation and reward well construction). Memory strength is assessed by calculating the percentage of time spent digging at the correct location and comparing it to random chance (which is 20% for 5 wells). Probe trials were interspersed to assess memory strength on sessions 11, 18, and 21, labelled PT1, PT2, and PT3, respectively. The percentage of dig time at the correct location was then compared across probe trials and against the chance level. As shown in *Fig1d*, the mean percentage of time spent digging at the correct location progressively increased from PT1 to PT3, indicating that memory strength improves with training (ANOVA, n=55, F-Value=5.5933, p= 0.006307). On PT3, the animals spent a significantly larger fraction of time at the correct location compared to random chance, while on both PT1 and PT2, the percentage of time spent digging at the correct location was at chance level (PT1: t-test, p =0.7684; PT2: 0.7899). This suggests that it took approximately 21 training sessions for the mice to learn the five flavours and their corresponding locations. The choice and composition of flavours, as well as the sand well locations relative to the intra-maze cues, were designed such that the animals learned all five flavours equally well (ANOVA, F-Value=1.0153; p= 0.40421; supplementary information1). From these results, we conclude that mice can successfully perform in an open event arena and learn 5 distinct flavour-place associations bound by the same context. We refer to this configuration of event arena as Map #1.

### Prior knowledge facilitates acquisition of new information of equal complexity *only when presented in relation to it*

In our task, the mice are required to learn the association between flavours and sand well locations, and by doing so, they also learn the spatial relationships between these locations within the arena. It is important to note that whenever the mice reach a particular location, the distances from the cues that define that location are reinforced irrespective of it being correct and rewarded or incorrect and hence not rewarded. This owing to the requirements of learning both correct as well as incorrect choices in discrimination training. Thus, we hypothesise that these implicitly acquired relations between locations along with explicit flavour-place pairs can act as *mental schema*. We then argued if this was to be true, then, new information when presented *in relation to prior knowledge* should result in faster acquisition (Tse et al. 2007). Specifically, we aimed to investigate whether schema-based learning could facilitate not just the acquisition of a few additional paired associations (as shown by Tse et al. 2007), but rather an entirely new event arena configuration (referred to as Map#2). The new configuration in Map#2 does not share any common features (such as intra-maze cues, flavours, or reward locations/sand wells) with Map#1. In this premise, we tested two questions: (i) can the animals learn this new map, and(ii) does the rate of learning depend on prior knowledge? There can be three possible outcomes:(a) there is no overlap between old and the new map memories. In that case, the new learning should resemble the learning of the initial map (Map#1); (b) the information content of Map#1 exceeds the animal’s cognitive load handling capacity, impairing new learning; or (c) old memories can assist in learning of the new map at a rapid rate.

To test this, we trained animals that had already learned Map#1 to acquire an additional set of five flavour-place associations using two different training routes: (i) **Solitary Learning:** Animals are trained on unique configuration of 5 novel flavour-place associations in a new context (i.e. Map#2) with no explicit reference to Map#1 (ii) **Relational Learning:** This approach takes advantage of the flexible nature of our behavioural paradigm, where Map#2 is introduced through gradual changes to Map#1.

We introduced a notation system to represent these modifications in the training paradigms. Each arena configuration is written as a set of alphanumeric characters, where the letters represent the intra-maze cues, and the numerals represent the unique flavour-location pairs. For example, animals that learned Map#1 (represented as AB12345) were trained on Map#2 (represented as CD67890) directly in the solitary learning group. In contrast, the relational learning group experienced a sequence of modifications: AB12367, AB12678, AB67890, AC67890, and finally, CD67890. After each modification (in relational learning grp), the mice underwent two training sessions on that configuration before the next change was introduced. The training paradigm and maps used are illustrated in the bottom panels of *Fig2a and Fig 2c*.

**Fig 2:**
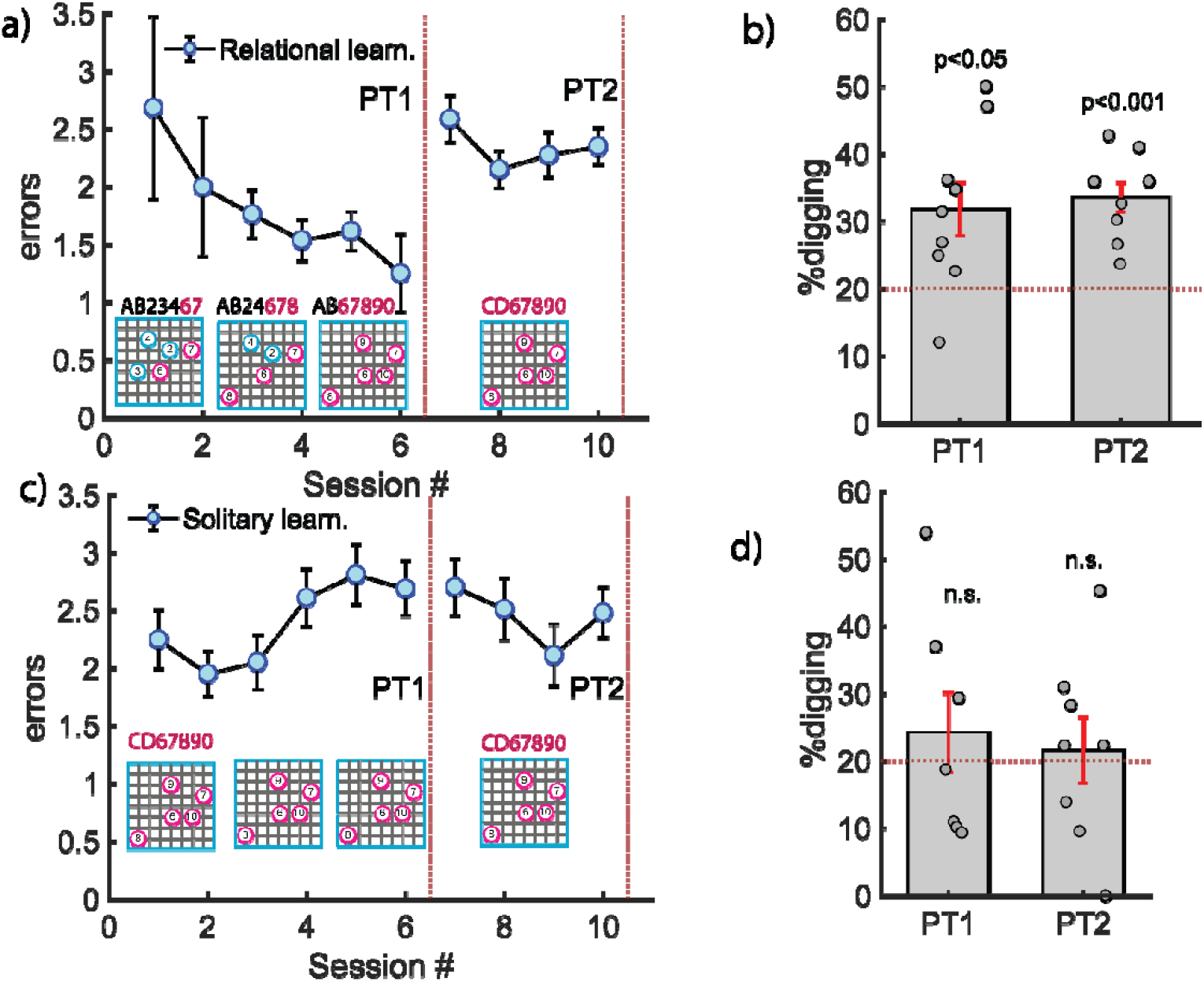
Acquisition of new flavour-location configuration (Map#2) by the two groups of mice that were trained using two different approaches. **(a)**Schematic representation of the training protocol followed to train two performance matched group of animals on new flavour-place configuration (Map2). For one group, Map1 was gradually morphed into Map2 (Relational Learning) while other group was trained on new map from the beginning (Solitary Learning). **(b)** Performance of relational learning and solitary learning group. The red dashed lines indicate the sessions corresponding to the probe trials. **(c)** Relational learning group performs significantly above chance during the probe trial for the new map with old cues (n=8, unpaired t-test, t-statistics=2.55077, p=0.03806) and with new cues (n=8, unpaired t-test, t-statistics=5.75607, p=0.000694074). The solitary learning group was unable to acquire the new map after 10 days. **(d)** Performance of solitary learning group during probe trials on day 6 (n=7, unpaired t-test, t-statistics=0.65832, p=0.53476) and day 10 (n=7, unpaired t-test, t-statistics= 0.72277, p=0.49702) of the training sessions.

The relational group progressed with faster acquisition of Map#2 compared to the solitary learning group (*Fig2a and Fig2c*). To assess memory strength after six training trials, we conducted a probe trial before changing the visual cues for the relational learning group. The results of this probe trial (*Fig2b PT1*) indicated that the animals in the relational learning group had successfully learned the new flavour-place associations (n=8, t-test, p=0.03806).

Following this, we modified the intra maze cues to eliminate any residual relationships from Map#1 thus resulting in the Map#2 configuration. It is important to note that, at this point, both the solitary and relational groups were exposed to the same Map#2 configuration (CD67890). Probe trial results after 10 training trials*Fig2b PT2* revealed that animals in the relational learning group spent 29.4% of their time digging at the correct locations during PT1 and 33.5% during PT2, indicating above-chance performance. In contrast, the solitary learning group animals spent 24.18% of their time digging at the correct locations during PT1 and 23.9% during PT2, with no significant difference from chance levels (*Fig2d* | n=7, t-test, p=0.535).

These results demonstrate that the animals acquire the new Map#2 faster only when presented in relation to the previously acquired Map#1 (in a graded manner). This suggests that mental schema not only facilitate faster acquisition of related information, but that new information, only when presented in relation to pre-existing schema (relational group) and not otherwise (solitary group), can enable acquisition of a complex multitude of new information.

### Graded Presentation is not Sufficient to Facilitate the Acquisition of Schema

We next considered the possibility that the enhanced cognitive load handling capacity (CLHC) observed in the relational group might have resulted from the presentation of novel information in smaller “chunks,” rather than being facilitated by prior learning, as we originally hypothesized. To test this, we conducted an experiment with another group of animals that were handled, habituated, and shaped alongside the relational group, but were directly trained on the flavour-place associations of Map#2 in a graded manner, without any prior training on Map#1. Note This approach mimics the training process of the relational learning group, with the key difference being the absence of any pre-existing schema for Map#1.

In this experimental design, the animals were first presented with 2 flavour-place pairs (ABxxx67) during the initial 2 trials. In the 3rd and 4th trials, an additional novel flavour-place pair (ABxx678) was introduced, and finally, all 5 flavours (AB67890) were presented during the 5th and 6th trials, similar to the relational learning group (*except no prior training for Map#1*). We hypothesized that if prior memory was not influencing learning, and the presentation of new information in small, manageable chunks was sufficient for rapid learning, then this control group should show faster and complete acquisition of all five flavour-place associations equivalent to relational learning group.

*Fig3a* shows the performance of these mice scored using the error measure as function of sessions. We observed a rapid initial learning, followed by a plateau and eventually a decline in performance. To assess memory strength, we conducted probe trials (similar to the relational and solitary experimental groups). The animals quickly learned to associate the first two flavour-location pairs. When probed for their associations, they performed exceptionally, retrieving the correct pairs with a performance significantly above chance (n=10, t-test, p=0.001) as shown in *Fig3b PT1*.

**Fig 3:**
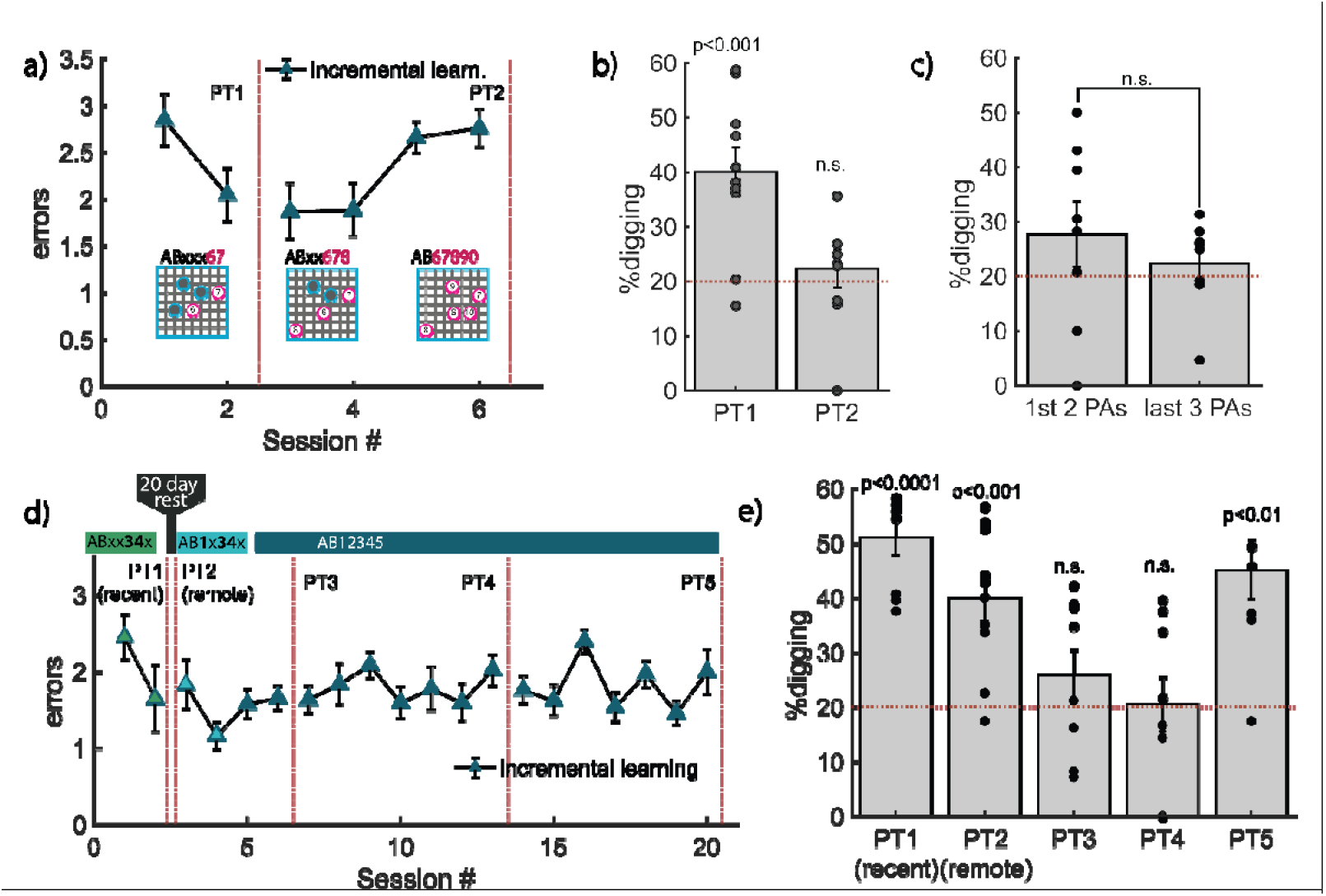
Acquisition of new flavour-place configuration (Map2) by incremental learning group mice. (a)Performance of the incremental learning group. The red dashed lines indicate site of probe trial. **(b)** The incremental learning group performs significantly above chance after training for two flavour-location associations for two days (n=10, unpaired t-test, t-statistics= 4.46688, p=0.00156), but are unable to acquire the other 3 flavour-location associations which were presented over the next 6 days (n=9, unpaired t-test, t-statistics=0.6517, p=0.53287). **(c)** When we assessed their performance on the second probe trial, we could clearly see that the new flavour-location associations have not been acquired. (d)(e)

However, as the training progressed and additional flavour-location pairs were introduced, the animals’ performance plateaued initially, then declined by session #6 (*Fig3a)*. After six sessions of training on this regime, a probe trial was conducted, and as shown in *Fig3b PT2*, the animals’ performance dropped to chance (n=9, t-test, p=0.532). This was surprising, given their strong performance during the first probe trial.

To investigate further, we divided the second probe trial data into two groups: (i) the 2 flavours initially presented (in Session #1 and #2), for which the animals had shown good memory during PT1, and (ii) the remaining 3 flavours. Since the training involved extensive randomization of flavour presentations and start box locations, both sets included all flavour-location combinations and were free from specific identity-based effects. Surprisingly, even for the flavours that the animals had successfully recalled during PT1, their performance at the second probe trial (*PT2*) was at chance (*Fig3c* | n=8, ANOVA, t-statistics=1.2957, p=0.237). This suggests that the introduction of additional flavours likely led to memory interference. Overall, these results indicate that while presenting new information in small chunks initially aided learning, it was not sufficient for encoding the full set of five associations. Furthermore, we conclude that training in small chunks, without the aid of pre-existing schemas, is more susceptible to memory interference.

This together with our results from relational and solitary training studies argues that the faster learning we see for a subset of flavour pair associates could be due to the “chunking” but they fail to encode any subsequent associations. Given these observations we infer that schema avoids catastrophic forgetting by helping the animal encode multitude of information while still preserving recently acquired prior knowledge.

### Generalisation does not reduce the CLHC

Generalization is thought to be a mechanism that reduces memory overload by forming representations that abstract and unify related memory events (Reinert et al. 2021). This process is believed to help the brain link different life experiences, thereby making it easier to learn similar information (Richards et al. 2014). In this study, we explored whether allowing sufficient time for a small portion of a complex memory to consolidate (Anagnostaras, Maren, and Fanselow 1999) can enhance cognitive load handling capacity for subsequent learning, without the need for a pre-existing mental schema.

We trained a group of animals on a 2 flavour-place association task over two days and tested their memory in probe trials administered on the 3rd day (*Fig3d*). As shown in *Fig3e, PT1*, the animals performed significantly above chance (n=8, t-test, p<0.001). After this, the animals rested in their home cage for 20 days. Following the rest period, we tested their remote retrieval after a single reminder session. The animals showed clear, above-chance digging at the correct location compared to incorrect locations (*Fig3e* | *PT2;* n=10, t-test, p<0.001).

Subsequently, we trained the animals using an incremental learning regime, where new flavour-place associations were introduced every 2 days. When tested for memory of these pairs after 4 days of training, the animals’ performance dropped to chance (*Fig3e* | *PT3;* n=9, t-test, p=0.287). This trend mirrored our observations in the Incremental learning group discussed earlier. These findings suggest that a long rest period for memory consolidation does not facilitate rapid learning if the initial learning set was not sufficiently complex or rich (only 2 paired associations).However, when training continued for a similar duration to our original Map#1 training, the animals exhibited above-chance performance for all 5 flavour-place pairs (*Fig3e* | *PT5;* n=8, t-test, p=0.008).

### Generation of Context Relevant Novel Responses

Having demonstrated that prior schema can facilitate the acquisition of new information, we sought to explore whether the same framework could help animals navigate novel scenarios derived from past experiences. This provides a foundation for considering “problem-solving behaviour” in rodent research. Specifically, we aimed to investigate whether the animals could deduce the correct location of the food pellet in a new, previously unknown schematic layout that is implicitly related to maps they had been exposed to earlier.

To test this, we exposed both the relational and solitary learning groups (which had been exposed to Map#1 and Map#2 equally) to a new configuration, Map#3 (CD12345). This map was entirely novel to the animals, but it could be derived from information previously learned in the earlier maps. We created Map#3 by taking the flavour-location pairs from Map#1 (xx12345) and placing them within the context of Map#2 (CDxxxxx), thus forming a hybrid configuration that the animals had not encountered before (*Fig4e*). This allowed us to test whether either group could reorient themselves based on new cues and dig at the correct food location, despite having no prior training on this map. The rationale behind this experimental design is that both groups had learned Map#1 and been equally exposed to Map#2 (10 sessions). To reduce anxiety from the novel map, we conducted probe trials for flavours #4 and #5 while training the mice on flavours 1 to 3 to prime them.

We found that animals in the relational learning group were able to solve the hybrid configuration by visiting the correct location significantly more often than the incorrect location (*Fig4b* | n = 8, Mann-Whitney U Test, U-value =14, critical value of U at p<0.05 = 15, Z score = 1.83787, p =0.03288). This trend of performance was absent in the animals that underwent solitary learning paradigm (*Fig4b*; n = 7, Mann-Whitney U Test, U-value =17, critical value of U at p<0.05 = 11, Z score = -0.89443, p =0.18673). We wanted to further substantiate if this performance is due to animals’ ability to relate the new visual cues with the Map#1 flavour-location associates or through some other unknown cues in the arena. This is critical to our hypothesis. One way to test this is through removing the visual cues and testing for their performance. When the visual cues were removed, both groups failed to perform the task. Specifically, the relational learning group visited the correct location significantly less often (n = 8, Mann-Whitney U Test, U-value =14, critical value of U at p<0.05 = 15, Z score = 1.83787, p =0.03288) and the solitary learning group showed no preference for the correct location, performing no differently than when the visual cues were present (n = 7, Mann-Whitney U Test, U-value =14, critical value of U at p<0.05 = 11, Z score = -1.27775, p =0.10027). These results suggest that the relational group is using the intra-maze landmarks from Map#2 to solve the task, while the solitary group, despite being trained on both Map#1 and Map#2 for the same duration, is unable to do so.

**Fig 4:**
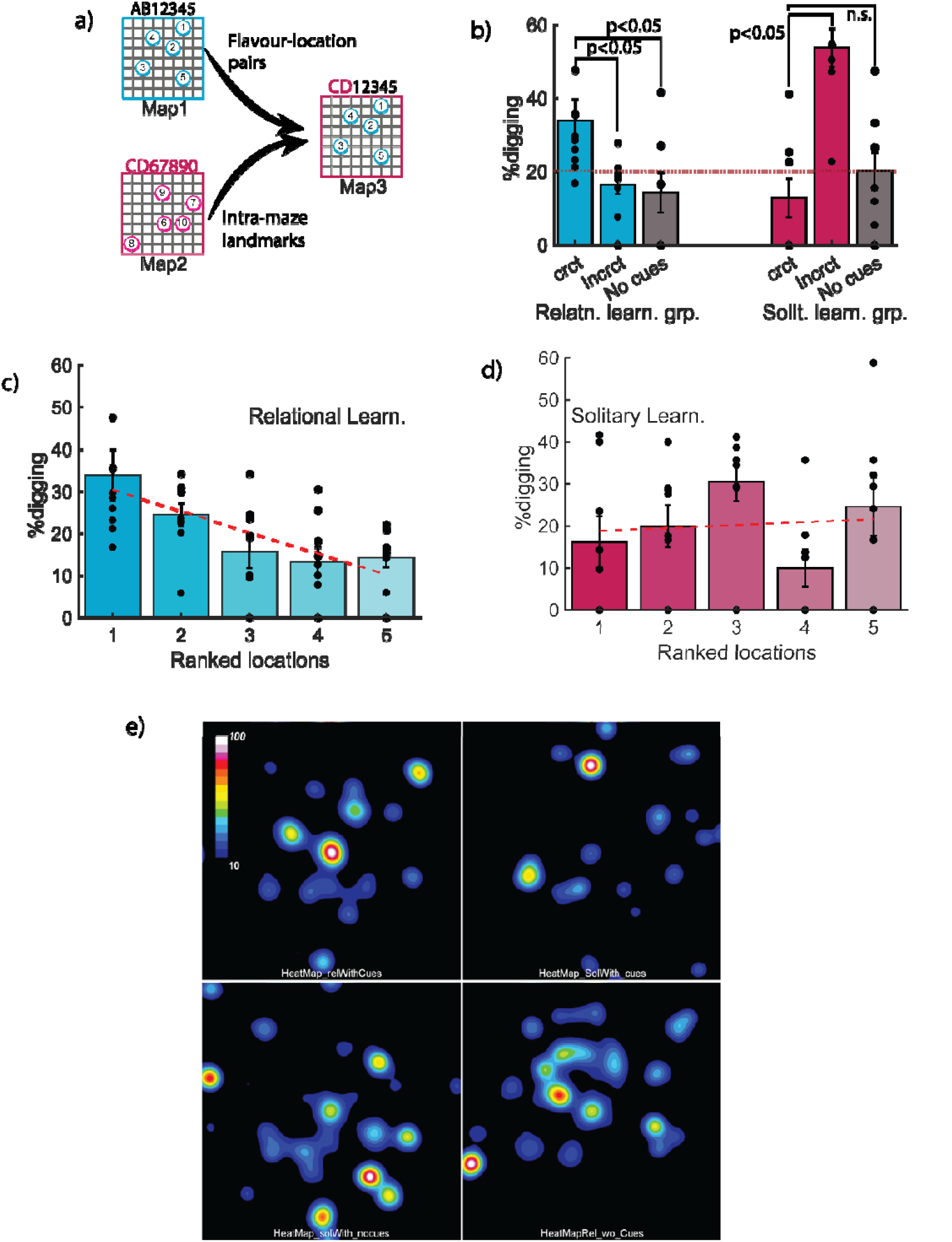
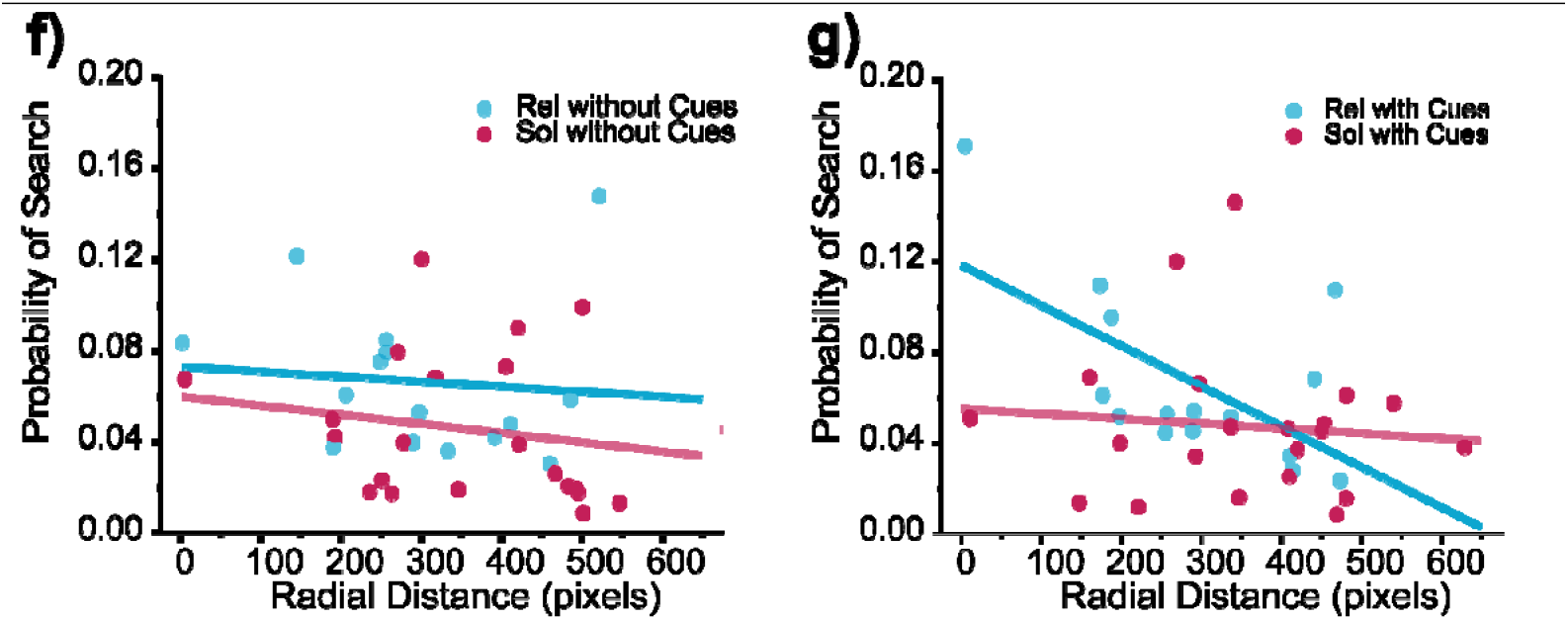
Performance of mice in a problem-solving task. **(a)** Schematic representation of the design for problem solving task. The favour-location associations were borrowed from Map#1 as it is the map learnt by both the group and visual cues were borrowed from Map#2. **(b)** The relational learning group mice visit the correct location significantly more often than the incorrect locations (n=8, Mann-Whitney U Test, U-value = 14, critical value of U at p<0.05 = 15, Z-Score = 1.83787, p=0.03288). Also, the difference in the animals’ performance with and without cue is significant (n=8, Mann-Whitney U Test, U-value = 14, critical value of U at p<0.05 = 15, Z-Score 1.83787, p=0.03288), suggesting that animals indeed use the visual cues to navigate and arrive at the correct location. For the solitary learning group of mice, there is no significant difference in their visit to correct and incorrect locations; either with (n=7, Mann-Whitney U Test, U-value = 17, critical value of U at p<0.05 = 11, Z-Score = -0.89443, p=0.18673) or without cue (n=7, Mann-Whitney U Test, U-value = 14, critical value of U at p<0.05 = 11, Z-Score = -1.27775, p=0 .10027). **(c)** Distance-wise ranking of flavour locations with respect to correct location in relational learning shows a downward trend suggesting that the animals follow neigh rest neighbour search pattern.; while **(d)** in case of solitary learning, they visit all the locations equally with no preference to the nearest neighbours. **(e)** The residence time heat map during problem solving trial. The individual trials of the mice are tracked and re-oriented such that the center of the image (512, 512) corresponds to the correct food pellet location. As can be seen from the map the group of animals that were trained in relation to prior memory (top left) not only search more at the correct location compared to animals trained in solitary group with (top right) and without visual cues (bottom left) and relative group without the visual cues (bottom right). they search in the vicinity of the correct location. Each spot represents location of one or more search center where food well was located. After reorienting the map to have the correct location at 512, 525, the residence times were added at each pixel and entire image was normalised for highest occupancy. **(f)** Shows the probability of search for reward as a function of distance from the correct location without landmarks in the arena vs **(g)** with landmarks in the arena for relational and solitary learning groups during problem solving.

We hypothesized that if the animals were truly using the cues to reorient themselves and were not simply dependent on the visual cues, their error patterns should reflect their process of reorienting in space. Specifically, as the animals learn to remap the locations, their digging behaviour should indicate their confidence in locating the reward. To test this, we measured the amount of time the animals spent digging at various locations, relative to their distance from the correct reward location. We ranked the reward locations based on their proximity to the correct location. Upon analysis, we observed a clear downward trend in the digging behaviour of the relational learning group (*Fig4c*), suggesting a convergent strategy in which the animals’ confidence in a location decreases as they move further away from the correct reward spot. However, in the solitary learning group, the animals did not exhibit such linear decrease in digging percentage as a function of spatial ranks (*Fig4d*). In fact, it seems that in this problem-solving task, the animals give equal weightage to every location (*Fig4d*). Next, we probed if such relationship is true when distance in measured in continuous space. We did this through generating residence time maps (heatmaps) that represent the time the animal spent in each spatial location with the correct location being the origin. The resulting heatmaps (*Fig4e*) for relational and solitary group clearly demonstrate the selective search exhibited by the animals (the bright centre spot) and progressive decline in the intensity of these search in space.

### Development of Physiologically Relevant ANN that Emulates Mental Schema Formation

Our findings from the experiments above suggest that future learning can benefit from pre-existing organised memory frameworks. The key factor in handling high cognitive load was minimizing interference, which is similar to the concept of catastrophic forgetting in artificial neural networks (ANNs)(French 1999; McCloskey and Cohen 1989). Hwu et. al. proposed a contrastive Hebbian learning ANN that is physiologically closer to neural communication as opposed to backpropagation based ANNs (Hwu and Krichmar 2020; Baldi and Pineda 1991; Movellan 1991; O’Reilly 1996; Ororbia and Mali 2019). Their approach works by restricting learning to a small set of nodes for specific stimuli, inspired by the concept of distinct engrams encoding various associations(Cai et al. 2016). We adapted this network architecture to replicate the behavioural results observed in our experiments. This experiment, underpinned by a theoretical framework, could guide the design of more focused intervention studies in the future. Details on the architecture and training protocols can be found in supplementary information 2.

In summary, the artificial neural network (ANN) alternates between two phases: (a) a fast-learning “hippocampus stream” that uses Hebbian updates for learning, and (b) a slow-learning “schema/cortical stream” that applies contrastive Hebbian updates, with hippocampus activation from the fast-learning phase serving as the teacher signal. The network architecture consists of multiple layers, with their use depending on the active stream and the learning phase of the network (*Fig5a*).

**Fig 5:**
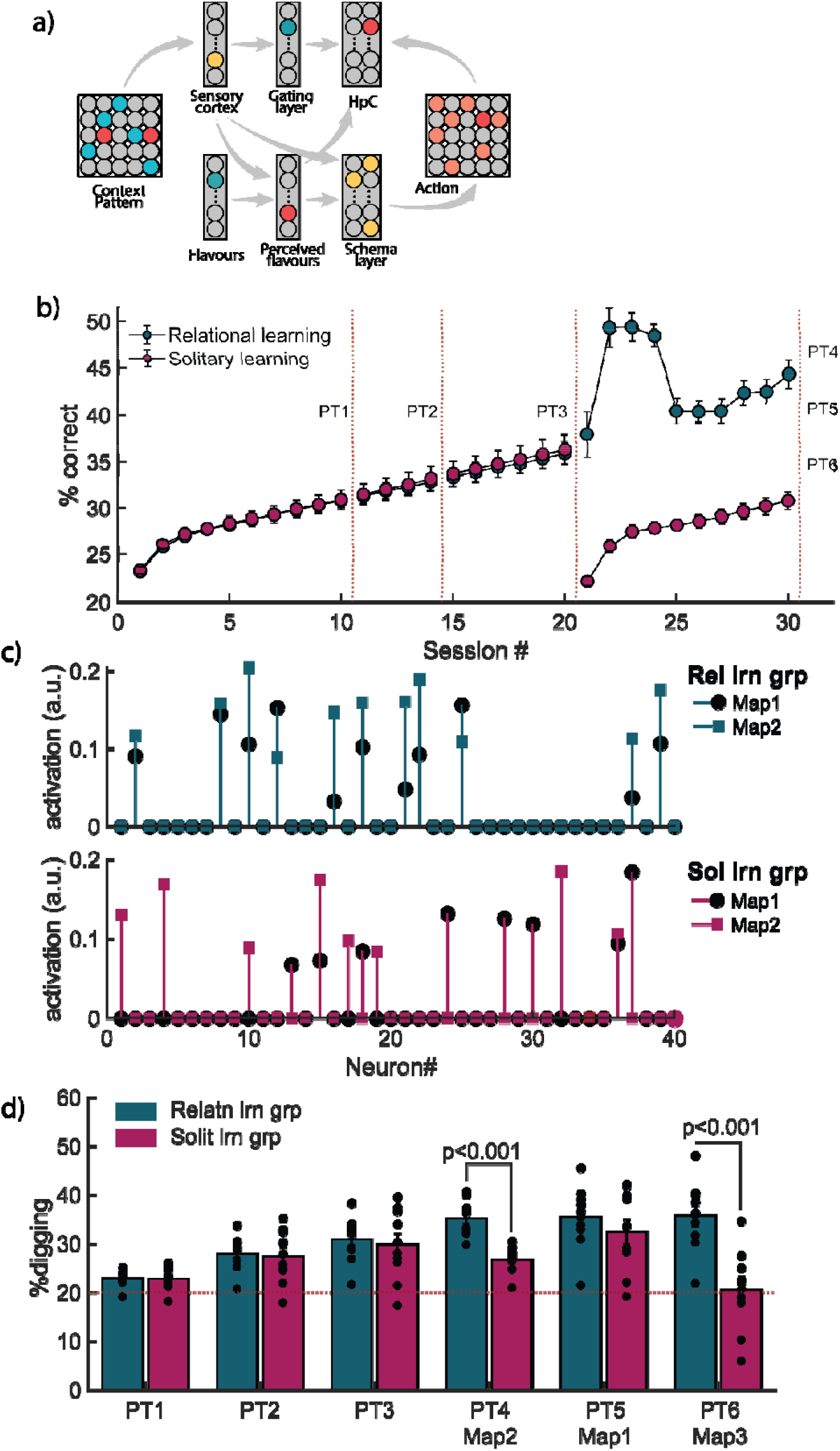
Performance of Artificial Neural network. **(a)** Architecture of the artificial neural network. Context patterns and flavours are the inputs for the system and the action is the output layer. The different routes of flow of information are shown with grey arrows. **(b)** Performance of the artificial neural network on 5 flavour-location associations till session 21. After this, half of the population was run through relational learning regime (cyan) and rest through the solitary learning (pink). **(c)** Relational learning group uses same nodes for learning both Map1 and Map2 while solitary learning group recruit different nodes for different maps. **(d)** The neural network learns Map#1 for 21 sessions during which we do not see any difference in the two populations of virtual animals (relational and solitary). After being trained for Map#2 as per the two different training regimes, we observe a significant difference in the performance of the two groups (PT4; Paired Sample t-Test; mean(rel) = 35.11364; mean(sol) = 26.65462; SD(rel)=3.62235; SD(sol)=2.7199; t Statistics = -4.8413; n = 10; p=9.19359 x 10^-4^) while their previous learning seems to be comparable (PT5 (relational); One Sample t-Test; mean(rel) = 35.36566; SD(rel)=6.451; t Statistics =7.53224; n = 10; p= 3.57012 x 10^-5^; PT5 (solitary); One Sample t-Test; mean(sol) = 32.35482; SD(sol)=8.27343; t Statistics =4.72227; n = 10; p=0.00109). These groups, when subjected to a problem-solving configuration, exhibited similar results as our mice studies wherein the relational learning group performed significantly better than the solitary learning group (PT6 (problem-solving); Paired Sample t-Test; mean(rel)= 35.68079; mean(sol)=20.51432; SD(rel)=7.03243; SD(sol)= 8.1939; t Statistics =-4.36769; n = 10; p=0.0018).

The network has three inputs: a context pattern (representing the task’s contextual configuration), a flavour, and the correct location associated with that flavour. The input context pattern (analogous to the learning map in our animal studies) is represented as a 2-dimensional grid of 25 elements (5 x 5). The reward locations, along with their relationships to other elements, are represented by an input value of 1, while landmarks, which have a significant influence on context representation, are represented by an activation value of 2. All other grid locations are set to 0. The flavour is represented by an 18-element array, with each active element corresponding to one of the 10 flavours used in the task. The action layer, which mirrors the context pattern’s dimensions, represents only the correct location for a given trial (as 1), with all other locations set to null. For more detailed description of the network architecture and training regime, please refer to supplementary information 2.

In our simulations, we began with 20 virtual mice divided into two groups: Relational learning and Solitary learning. We allowed the animals to learn the Map1 configuration for 21 days, similar to our animal study, with probe trials conducted after each session. We then subjected them to two different training protocols as described above. We observed that the network used the same nodes (during the CHL cortical stream) for the new map configuration in case of relational learning and was able to assimilate the new information much faster (*Fig5c*). At the end of 10 sessions (to achieve Map2), similar to our results from earlier experiments, the virtual animals participating in solitary learning regime performed poorly as compared to the relational learning group (*Fig5d PT4*; n=10; paired t-test; n=10; t-statistics: -4.8413; p<0.0001). Furthermore, when tested on the problem-solving configuration (Map3, combining flavour-location pairs from Map1 with intra-maze cues from Map2), the relational learning group outperformed the solitary learning group, similar to our behavioural results (*Fig5d PT6*). The Relational learning group was superior in performance as compared to solitary learning group (n =10; paired t-test; t-statistics: -4.36769; p=0.0018). A crucial factor in eliciting these results was the cross-inhibition between the schema layer and the fast-learning stream. Simulations without this cross-connection failed to produce correct responses during problem-solving. This is one of the key differences between our proposed ANN and previous models (Hwu and Krichmar 2020). Although our architecture is nearly identical to that of Hwu et al. (2020), we incorporated an additional layer of abstraction for flavours and context to ensure that the network did not treat the problem-solving map as a novel map. These encouraging results suggest that our approach could be used to design more targeted intervention studies in the future, exploring the role of engrams in learning and memory for complex tasks.

## Discussion

We introduce a novel event arena-based behavioural paradigm in rodents to investigate the role of prior knowledge and the adoption of different learning strategies in the acquisition of new information. Our findings demonstrate that learning in the relation to existing knowledge prevents catastrophic interference and aids in problem-solving tasks in mice. Previous research has shown that heavy cognitive load can hinder new learning (Kolibius, Born, and Feld 2020; Feld, Weis, and Born 2016). In contrast, our experiments reveal that animals are able to learn a complex open maze task (with more than five paired associates, as opposed to one or two) only when it is framed in relation to their prior knowledge.

While various aspects of mental schemas have been demonstrated in rats (Tse et al. 2007; Alonso et al. 2021; Wang and Morris 2010), here we show for the first time that mental schemas facilitate the acquisition of a completely new map of equal content and complexity. This is distinctly different from acquiring just a few new paired associates. We demonstrate this through our incremental learning group, where animals were able to perfectly acquire 2 flavour-place associations after just two training sessions, without any prior training. However, they lost these memories once additional flavour-place associations were introduced. In contrast, when the same training scheme is applied to animals with prior learning, they not only acquire the associations more quickly but also retain all 5 paired associates. We interpret this effect as evidence that mental schemas enhance an animal’s ability to handle cognitive load.

In demonstrating the above, we also establish an event arena-based task to investigate flavour-place associations. Our setup consisted of a 1.4 m x 1.4 m arena with five reward wells and two intra-maze cues. Each well was associated with a uniquely flavoured food pellet, and the animal was presented with the corresponding flavour pellet outside the arena before each trial. Throughout the experiment, we used probe trials (cued but unrewarded) to show that the mice were able to learn the maze and successfully associate the cued flavour with its respective location in the arena by utilizing the intra-maze cues.

Once the animals had acquired the initial flavour-location associations with respect to the intra-maze cues (Map1), we tested the utility of the “*mental schema* “ acquired in Map1by gradually modifying Map1 to create a completely new configuration, Map2 (referred to as relational learning). As a control, we also ran a separate group that, after learning Map1, was directly transferred to Map2 (referred to as solitary learning). This allowed us to compare learning outcomes with and without referencing acquired memories. We found that the relational learning group rapidly acquired Map2, while the solitary learning group remained at chance level for the same duration of training on the new map.

Since the relational learning group was presented with only a few changes per trial, their rapid learning behaviour may have resulted from chunking (Burtis 1982; Gobet and Clarkson 2004). To investigate role of chunking, we designed the *“Incremental learning group*”. Please note that this is a naïve experimental group that was never trained on Map1. We began by giving the animals just 2 associations in the arena, which they learnt rapidly. Following this, we introduced 3 additional unique associations (while keeping the old associations intact). After training the animals for two days on the final configuration (similar to the training procedure for the relational learning group), we conducted another probe trial. A careful analysis of the probe trial revealed that not only did the animals fail to learn all 5 flavour-place associations, but they also appeared to have forgotten the 2 flavour-place associations they had previously learned. These results suggest catastrophic forgetting due to memory interference (Navarro Lobato et al. 2023).

Combining the results from the relational, solitary, and incremental learning groups, we argue that having a rich memory association network (*“mental schema”*) facilitates faster acquisition, enhances high genitive load handing capacity, and prevents memory interference from similar associations acquired within close temporal windows.

Since the incremental learning group successfully learned the initial two association pairs quickly, we were curious to test whether this, combined with system consolidation, could produce the effects observed in the relational learning group. We trained a group of animals on 2 flavour-place associations and then gave them a long rest period (20 days), a duration shown in rodent literature to be associated with system consolidation(Li et al. 2023; Bonaccorsi et al. 2013; Terranova et al. 2023; Frankland and Bontempi 2005; Kim and Fanselow 1992). After the rest period, the animals were returned to the arena, given reminder training, and tested on their memory of the task. The animals performed above chance. We then proceeded with adding new flavour-place associations (similar to the incremental and relational learning paradigms) and observed results similar to those in the incremental learning group. Not only did the animals fail to acquire the new flavour-place associations, but it appeared that their previously learned associations were also compromised. This led us to conclude that the effects seen in the relational learning group cannot be solely explained by system consolidation or chunking but must instead involve a higher-order cognitive framework, such as a mental schema. While we acknowledge that further intervention studies will be needed to fully understand the role of system consolidation in this group, we took steps to rule out the possibility that these animals were simply incapable of learning a complex arena setup. We continued training them on the Map1 configurations, and after 20 sessions, the animals successfully acquired the new map and performed above chance during probe trials.

Mental schemas have been proposed as a framework for knowledge, playing a key role in decision-making and problem-solving (van Kesteren et al. 2012; Sweller 1988; Brewer and Treyens 1981; Lewis and Anderson 1985). Given that our results suggest mice may use a similar abstract structure, we aimed to investigate whether our two groups—one explicitly trained to relate new information to prior knowledge (relational learning group) and the other not given this explicit connection (solitary learning group)—would show differences in problem-solving scenarios. We created a hybrid map by merging visual cues from Map2 with flavour-place associations from Map1. We found that the relational learning group performed significantly better in solving the hybrid map compared to the solitary learning group. Furthermore, when intra-maze cues were removed, the performance of both groups dropped to chance level. This suggests that the animals were relying on the visual cues from the other map to navigate the hybrid configuration.

It has been postulated and well-documented that distinct engrams are formed for perceptually and cognitively different types of learning, while similar engrams encode overlapping cues that occur close in time. Drawing from this literature, Hwu et. al. (Hwu and Krichmar 2020) developed a contrastive Hebbian learning-based artificial neural network designed to replicate the behavioural findings of Tse et al.(Tse et al. 2007) on mental schemas and memory consolidation. The main advantages of this architecture and learning mode are twofold: First, the network adjusts for errors without backpropagation, making it more biologically plausible than traditional backpropagation methods. Second, it offers a solution to the problem of catastrophic forgetting by ensuring that distinct engrams encode different or novel information. We applied this network, with some modifications, to our dataset. Our results show that we could engage the same artificial engram for the relational learning group, while the engram switched for the solitary learning group. Furthermore, we demonstrated that this network could perform problem-solving tasks using a hybrid map, similar to the behaviour of the relational learning group in our experiments. This is particularly promising as it aligns with the current understanding of how engrams encode new information and sets the stage for future histological and intervention studies. The complexity and extended nature of the task made it difficult to investigate the neurological underpinnings through lesions. Future studies could explore the brain regions involved in this behaviour through passive observational techniques that use minimally invasive procedures.

## Materials and methods

### Animals

We used adult (5–6-month-old) C57BL/6J male mice from this study acquired from the Central Animal Facility, Indian Institute for Science. The animals were housed in pairs on a 12-h light/dark cycle. Before starting the experiment, one of the animals from each cage was briefly anaesthetized using 2% isoflurane and ear marked by making a small wedge cut on right ear. This helps us to identify them during the experimental run. The study is performed in accordance with the animal protocol approved by the Institute Animal Ethics Committee, Indian Institute for Science, Bengaluru.

### Event Arena

The main design of the event arena was inspired form the study by Tse et. al (Science, 2007). We adapted their design and made significantly design changes to adapt the setup to our experiments in rodents. Event arena consists of a 104 x 104 x 15 cm roofless cuboid box with 4 entry/starting boxes in the middle of each side. The floor of the arena is covered with fresh sand up to a height of 1-2 cm. The arena is held on metal frame and enclosed with an outer larger box to avoid animals using any external cues from the room. Animals’ behaviour was recorded using a HD camera mounted on the roof of the external box. The reward wells (digging sites for animal to acquire reward) were placed in arena to avoid any rotational symmetry. The position and number of these reward wells and intra-maze landmarks depends on the experimental run.

### Habituation

The mice are handled twice daily on top of their home cage for (6–10 days) to wean off any anxious escape behaviours and acclimate them to human handling. Following this, the animals are allowed to explore the arena with food pellets randomly scattered throughout the arena. Habituation is deemed complete once the animals stop wall hugging and freely exploring the centre of the arena.

### Food restriction

The food restriction was implemented in a gradual manner with restricting them to two pellets per mouse per day for 1^st^ week and then tapering it down to 1 pellet per mouse per day for next few days. Once the experimenters are sure that the animals have acclimated to the regime, the animals were started on shaping and training protocols. On experiment days, the animals are given only 1 pellet per mouse outside of the experimental arena while on off days they are fed 2 pellets per day. Experimenter kept a close watch on the animal behaviour and if any animal exhibited lethargy and hunching, the animal was given more food and excluded from the experimental run till recovery.

### Shaping

Our experimental design relies on the animals to dig the favoured pellet from under the sand. To induce digging behaviour in the animals, we perform shaping. To begin with, the animals are presented with food reward on the top of sand wells. Note that there is no flavour present in the food pellets for shaping trials. The food pellet is then gradually lowered into the sand well on subsequent shaping sessions. The shaping is considered completed when the food pellet reaches the bottom of the sand well (3 cm deep) and the mice are able to retrieve the pellets.

### Flavour pellet

The flavoured food pellets were prepared inhouse by mixing flavours to ground up rodent feed (NutriLab Rodent Feed, Provimi) and then extruding the paste into pellets of desired sizes. The pellets were allowed to dry in an oven for 4 hours. All the procedures were performed in a semi-sterile environment. Fresh pellets were made every week. The pellets were labelled and stored in air-tight containers and stored in dry and dark conditions. The various flavours used for the experiment are as follows:

| S. No | Flavour | Supplier | Concentration (w/w) |
| --- | --- | --- | --- |
| 1. | Basil | Snapin, Duval enterprises Pvt. Ltd. | 1% |
| 2. | Cassia | Arvind Trading Company | 0.50% |
| 3. | Cocoa | WeikField, Weikfield foods Pvt. Ltd. | 1% |
| 4. | Cumin | Everest, S. Narendrakumar & Co. | 1% |
| 5. | Dry Mango | Everest, S. Narendrakumar & Co. | 1% |
| 6. | Elaichi | Arvind Trading Company | 1% |
| 7. | Fenugreek | Eastern, Eastern Condiments Pvt. Ltd. | 1% |
| 8. | Ginger | Snapin, Duval enterprises Pvt. Ltd. | 1% |
| 9. | Rosewhite | Bush, International Flavors and fragrances India Pvt. Ltd. | 1% (v/w) |
| 10. | Thyme | Keya, Keya Foods International Pvt. Ltd. | 0.50% |

### Training protocol

Every experimental run is called a session which comprises of 5 individual trials with non-repeating pseudo-randomly assigned flavour-place pair per trial. Note that the relationship between a flavour and its corresponding location in the arena never changes over the course of the experiment while the sequence in which the pairs are presented and site of entry to arena is randomized per session per mice. At the start of a trial, the animal is presented with the cue flavoured pellet and allowed to consume the pellet for about 20 sec. Following this, the animal is dropped at the randomly assigned site of entry and the session begins as soon as the animal leaves the start box and enters the arena. The mice are given 120 seconds to find the correct pellet and if the animal fails, the experimenters suggest the correct location and give then an extra 30 seconds to retrieve the pellets. After the mouse has dug and retrieved the pellet, it has 10–15 s to consume the food reward before it is returned to the home cage. Once the trial is over, next correct reward well is baited and all the reward wells are replaced with fresh ones. The sand is sprayed with 70% alcohol, sand is shuffled and flattened to remove any odour or visual track cues.

In general, the experiments are run such that there is at least 45 min delay between each trial for any given mouse.

### Cued probe trials

We test the strength of the memory using cued probe trials wherein the animal is cued while the correct location is devoid of any food reward. This ensures that there are no olfactory cues helping the animal arrive at the correct location. If the animal is certain of the correct location, they revisit the location over and over and spend significantly more time digging at the location despite no food pellet being there. To avoid extinction, the probe trials are run for only 60 sec and the experiments rewards the animal at the correct location with the correct flavoured pellet at the end of the probe trial. Performance during probe trial is measured as the percentage of digging time at the correct location over the total digging time for the trial.

### Artificial Neural Network training

The neural network was custom written in MATLAB 2014 and was run on a i5-32GB-CPU. The data input for various learning setups is provided in supplementary information 2.

## Data and code sharing

Data and code for the study can be shared after acceptance of the manuscript upon a reasonable request to the authors.

