## Supplemental Information for "Mental Schema Reduces Cognitive Load and Facilitates Emergence of Novel Responses in Mice and Artificial Neural Networks"

### Supplementary information and figures:

*Supplementary information 1:* ***Performance of animals split by flavour during PT3 Map1***

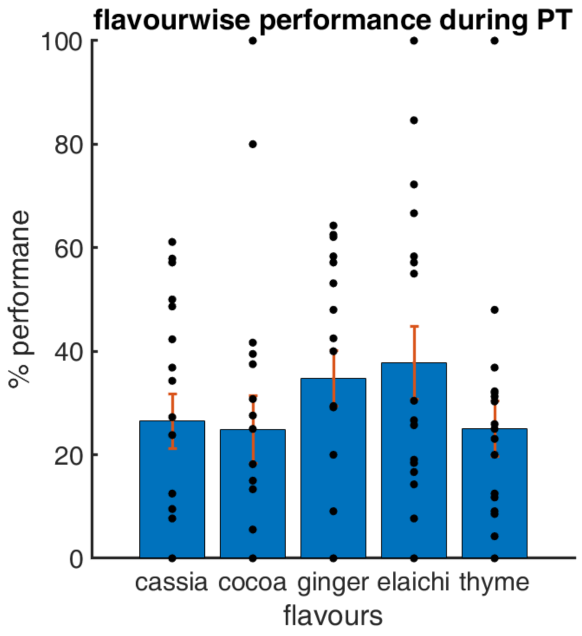


*Supplementary fig 1: flavour-wise performance of mice on PT3 for Map1*

| | SumOfSquares | DF | MeanSquares | F | pValue |
|---|---|---|---|---|---|
| **Flavours** | 2619.9 | 4 | 654.97 | 1.0153 | 0.40421 |
| **Error** | 54834 | 85 | 645.11 | | |
| **Total** | 57454 | 89 | | | |

*Supplementary information 2:* ***ANN architecture***

#### *a)* ANN architecture:

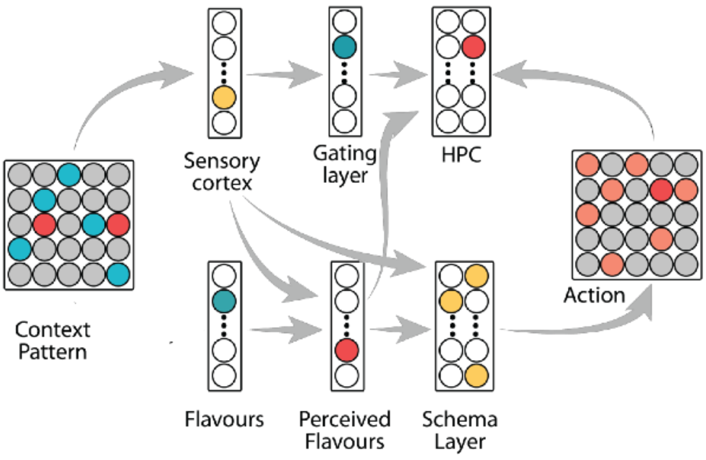


*Supplementary fig 2: ANN architecture.*

#### *b)* Initialization of different layers:

**The various layers are initialized as follows:**

**I. Context Pattern:**

| | |
|---|---|
| **Schema A** | **[0,0,1,0,0;0,1,0,0,0;0,2,0,1,2;1,0,0,0,0;0,0,0,0,1]** |
| **Map1_1 (Relational learn.)** | **[0,1,0,0,0;0,0,1,0,0;0,2,0,1,2;1,0,0,0,0;0,0,0,0,1];** |
| **Map1_2 (Relational learn.)** | **[0,1,0,0,0;0,0,1,0,0;0,2,0,1,2;1,0,0,0,0;1,0,0,0,0];** |
| **Map2 (Relational learn.)** | **[0,1,0,0,0;0,0,1,0,0;1,2,0,0,2;0,0,0,1,0;1,0,0,0,0];** |
| **Map2 (relational learning with new landmark position)** | **[0,1,0,2,0;0,0,1,0,0;1,0,0,0,0;0,0,0,1,0;1,0,2,0,0];** |
| **Map2 (Solitary learn)** | **[0,1,0,2,0;0,0,1,0,0;1,0,0,0,0;0,0,0,1,0;1,0,2,0,0];** |

**II. Cue sets:**

| | |
|---|---|
| **Schema A** | **[1,0,0,0,0,0,0,0,0,0,0,0,0,0,0,0,0,0;...<br>0,1,0,0,0,0,0,0,0,0,0,0,0,0,0,0,0,0;...<br>0,0,1,0,0,0,0,0,0,0,0,0,0,0,0,0,0,0;...<br>0,0,0,1,0,0,0,0,0,0,0,0,0,0,0,0,0,0;...<br>0,0,0,0,1,0,0,0,0,0,0,0,0,0,0,0,0,0;];** |
| **Map1_1 (Relational learn.)** | **[0,0,0,0,0,1,0,0,0,0,0,0,0,0,0,0,0,0;...<br>0,0,0,0,0,0,1,0,0,0,0,0,0,0,0,0,0,0;...<br>0,0,1,0,0,0,0,0,0,0,0,0,0,0,0,0,0,0;...<br>0,0,0,1,0,0,0,0,0,0,0,0,0,0,0,0,0,0;...** |

| | |
|---|---|
| | 0,0,0,0,1,0,0,0,0,0,0,0,0,0,0,0,0,0;]; |
| **Map1_2**<br>**(Relational learn.)** | [0,0,0,0,0,1,0,0,0,0,0,0,0,0,0,0,0,0;...<br>0,0,0,0,0,0,1,0,0,0,0,0,0,0,0,0,0,0;...<br>0,0,1,0,0,0,0,0,0,0,0,0,0,0,0,0,0,0;...<br>0,0,0,1,0,0,0,0,0,0,0,0,0,0,0,0,0,0;...<br>0,0,0,0,0,0,0,0,0,1,0,0,0,0,0,0,0,0;]; |
| **Map2**<br>**(Relational learn.)** | [0,0,0,0,0,1,0,0,0,0,0,0,0,0,0,0,0,0;...<br>0,0,0,0,0,0,1,0,0,0,0,0,0,0,0,0,0,0;...<br>0,0,0,0,0,0,0,1,0,0,0,0,0,0,0,0,0,0;...<br>0,0,0,0,0,0,0,0,1,0,0,0,0,0,0,0,0,0;...<br>0,0,0,0,0,0,0,0,0,1,0,0,0,0,0,0,0,0;]; |
| **Map2**<br>**(Relational learning with new landmark position)** | [0,0,0,0,0,1,0,0,0,0,0,0,0,0,0,0,0,0;...<br>0,0,0,0,0,0,1,0,0,0,0,0,0,0,0,0,0,0;...<br>0,0,0,0,0,0,0,1,0,0,0,0,0,0,0,0,0,0;...<br>0,0,0,0,0,0,0,0,1,0,0,0,0,0,0,0,0,0;...<br>0,0,0,0,0,0,0,0,0,1,0,0,0,0,0,0,0,0;]; |
| **Map2**<br>**(Solitary learn)** | [0,0,0,0,0,1,0,0,0,0,0,0,0,0,0,0,0,0;...<br>0,0,0,0,0,0,1,0,0,0,0,0,0,0,0,0,0,0;...<br>0,0,0,0,0,0,0,1,0,0,0,0,0,0,0,0,0,0;...<br>0,0,0,0,0,0,0,0,1,0,0,0,0,0,0,0,0,0;...<br>0,0,0,0,0,0,0,0,0,1,0,0,0,0,0,0,0,0;]; |

**III. Action/output layer:**

| | |
|---|---|
| **Schema A** | [0,0,1,0,0,0,0,0,0,0,0,0,0,0,0,0,0,0,0,0,0,0,0,0,0;...<br>0,0,0,0,0,0,1,0,0,0,0,0,0,0,0,0,0,0,0,0,0,0,0,0,0;...<br>0,0,0,0,0,0,0,0,0,0,0,0,0,1,0,0,0,0,0,0,0,0,0,0,0;...<br>0,0,0,0,0,0,0,0,0,0,0,0,0,0,0,1,0,0,0,0,0,0,0,0,0;...<br>0,0,0,0,0,0,0,0,0,0,0,0,0,0,0,0,0,0,0,0,0,0,0,0,1;]; |
| **Map1_1**<br>**(Relational learn.)** | [0,1,0,0,0,0,0,0,0,0,0,0,0,0,0,0,0,0,0,0,0,0,0,0,0;...<br>0,0,0,0,0,0,0,1,0,0,0,0,0,0,0,0,0,0,0,0,0,0,0,0,0;...<br>0,0,0,0,0,0,0,0,0,0,0,0,0,1,0,0,0,0,0,0,0,0,0,0,0;...<br>0,0,0,0,0,0,0,0,0,0,0,0,0,0,0,1,0,0,0,0,0,0,0,0,0;...<br>0,0,0,0,0,0,0,0,0,0,0,0,0,0,0,0,0,0,0,0,0,0,0,0,1;]; |
| **Map1_2**<br>**(Relational learn.)** | [0,1,0,0,0,0,0,0,0,0,0,0,0,0,0,0,0,0,0,0,0,0,0,0,0;...<br>0,0,0,0,0,0,0,1,0,0,0,0,0,0,0,0,0,0,0,0,0,0,0,0,0;...<br>0,0,0,0,0,0,0,0,0,0,0,0,0,1,0,0,0,0,0,0,0,0,0,0,0;...<br>0,0,0,0,0,0,0,0,0,0,0,0,0,0,0,1,0,0,0,0,0,0,0,0,0;...<br>0,0,0,0,0,0,0,0,0,0,0,0,0,0,0,0,0,0,0,0,1,0,0,0,0;]; |
| **Map2**<br>**(Relational learn.)** | [0,1,0,0,0,0,0,0,0,0,0,0,0,0,0,0,0,0,0,0,0,0,0,0,0;...<br>0,0,0,0,0,0,0,1,0,0,0,0,0,0,0,0,0,0,0,0,0,0,0,0,0;...<br>0,0,0,0,0,0,0,0,0,0,1,0,0,0,0,0,0,0,0,0,0,0,0,0,0;...<br>0,0,0,0,0,0,0,0,0,0,0,0,0,0,0,0,0,0,1,0,0,0,0,0,0;...<br>0,0,0,0,0,0,0,0,0,0,0,0,0,0,0,0,0,0,0,0,1,0,0,0,0;]; |
| **Map2**<br>**(Relational learning with new landmark position)** | [0,1,0,0,0,0,0,0,0,0,0,0,0,0,0,0,0,0,0,0,0,0,0,0,0;...<br>0,0,0,0,0,0,0,1,0,0,0,0,0,0,0,0,0,0,0,0,0,0,0,0,0;...<br>0,0,0,0,0,0,0,0,0,0,1,0,0,0,0,0,0,0,0,0,0,0,0,0,0;...<br>0,0,0,0,0,0,0,0,0,0,0,0,0,0,0,0,0,0,1,0,0,0,0,0,0;...<br>0,0,0,0,0,0,0,0,0,0,0,0,0,0,0,0,0,0,0,0,1,0,0,0,0;]; |
| **Map2** | [0,1,0,0,0,0,0,0,0,0,0,0,0,0,0,0,0,0,0,0,0,0,0,0,0;... |

| | |
|---|---|
| **(Solitary learn)** | **0,0,0,0,0,0,0,1,0,0,0,0,0,0,0,0,0,0,0,0,0,0,0,0,0;...<br>0,0,0,0,0,0,0,0,0,0,1,0,0,0,0,0,0,0,0,0,0,0,0,0,0;...<br>0,0,0,0,0,0,0,0,0,0,0,0,0,0,0,0,0,0,1,0,0,0,0,0,0;...<br>0,0,0,0,0,0,0,0,0,0,0,0,0,0,0,0,0,0,0,0,1,0,0,0,0;];** |

**c) Initial weights for each layer to contribute to other layers is randomized.**

#### *d)* Training protocol:

- The dHPC layer receives three inputs during the hippocampal learning stream: the contextual pattern (via sensory layer and gating layer); the perceived flavour (the perceived flavour is the product of the sensory layer activation and the flavour layer. This is done to account for the effect of the context on the perceived information); and the corresponding correct location for the task from the action layer (*supplementary fig. 3*).

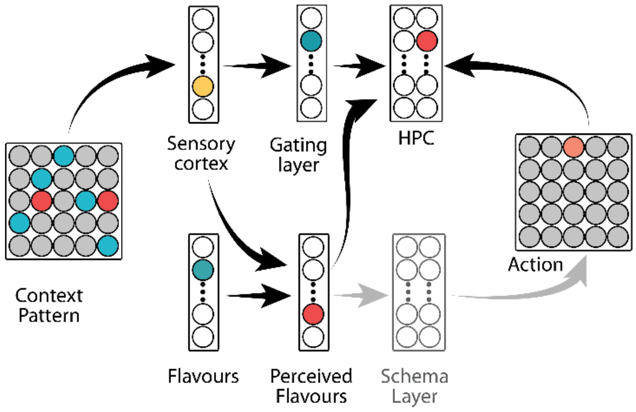


*Supplemetary fig 3: Hippocampal training architecture*

- The network goes through multiple epochs, and once the activity of each layer converges, the network enters the cortical learning stream where the weight updates happen via contrastive Hebbian Learning.
- The contrastive Hebbian learning depends on the difference in the weights in the two phases of any network i.e. a free phase and a teacher phase (*supplementary fig 4*).

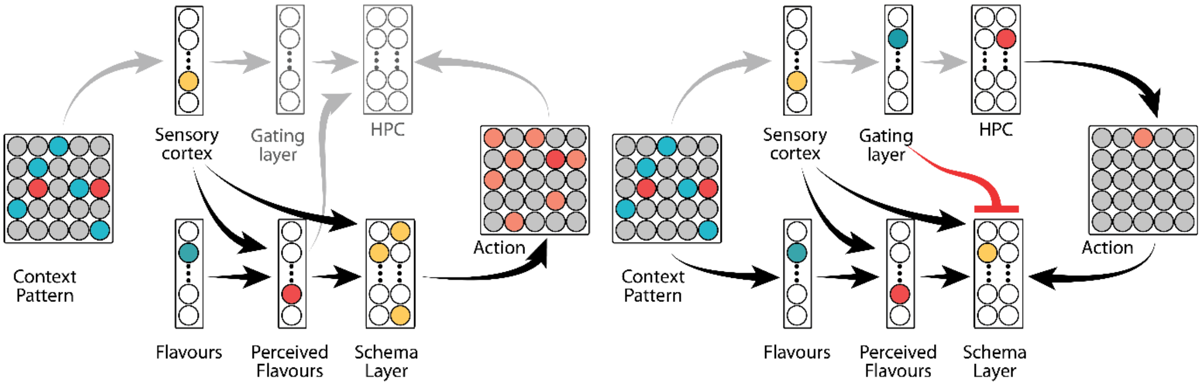


*Supplementary fig 4: Left: free phase; Right: teacher phase*

- The cortical stream begins with the free phase where the schema layer receives two inputs: the context pattern and flavour (via same layers as the hippocampal stream) and is allowed to produce a correct location in the action layer. This is followed by a teacher phase where the dHPC active neuron (from the previous dHPC learning phase for a particular flavour-location trial) acts as a teacher for the correct location to the schema layer. During this phase, the schema layer receives 3 inputs (context pattern, flavours and correct location). Additionally, in this phase, the nodes in schema layers are gated in a manner where only a few nodes can participate in this learning. This is to emulate the fact that not all the cells participate in every learning and only a subset of neurons encode the incoming information. This is achieved by using a gating layer to project negative inhibitory weights to the schema layer with a few elements of the inhibitory matrix set to zero (not affecting the downstream layer).

- The performance of the cortical stream in the two phases is used to implement contrastive Hebbian weight updates for every layer. Another feature of this ANN is that at each trial it calculates the familiarity of the context pattern and modulates the number of learning epochs (see supplementary information). This implementation allows us to detect schema with consistent and inconsistent information.
